# Deep Learning-based Framework for Mycobacterium Tuberculosis Bacterial Growth Detection for Antimicrobial Susceptibility Testing

**DOI:** 10.1101/2025.02.14.638231

**Authors:** Hoang-Anh T Vo, Sang Nguyen, Ai-Quynh T Tran, Han Nguyen, Hai Ho Bich, Philip W Fowler, Timothy M Walker, Thuy Thi Nguyen

## Abstract

Tuberculosis (TB) kills more people annually than any other pathogen. Resistance is an ever-increasing global problem, not least because diagnostics remain challenging and access limited. 96-well broth microdilution plates offer one approach to high-throughput phenotypic testing, but they can be challenging to read. Automated Mycobacterial Growth Detection Algorithm (AMyGDA) is a software package that uses image processing techniques to read plates, but struggles with plates that exhibit low growth or images of low quality. We developed a new framework, TMAS (TB Microbial Analysis System), which leverages state-of-the-art deep learning models to detect *M. tuberculosis* growth in images of 96-well microtiter plates. TMAS is designed to measure Minimum Inhibitory Concentrations (MICs) rapidly and accurately while differentiating between true bacterial growth and artefacts. Using 4,018 plate images from the CRyPTIC (Comprehensive Resistance Prediction for Tuberculosis: An International Consortium) dataset to train models and refine the framework, TMAS achieved an essential agreement of 98.8%. TMAS offers a reliable, automated and complementary evaluation to support expert interpretation, potentially improving accuracy and efficiency in tuberculosis drug susceptibility testing (DST).

**Author summary:** Tuberculosis (TB) is one of the world’s leading causes of death from infectious diseases, with drug resistance making treatment increasingly difficult. Accurate and timely drug susceptibility testing (DST) is essential to determine which antibiotics remain effective against a given TB strain. A widely used DST method involves 96-well broth microdilution plates, where bacteria are exposed to different antibiotic concentrations to determine the minimum inhibitory concentration (MIC), the lowest concentration that fully prevents bacterial growth. However, manually interpreting these plates is time-consuming and prone to human error, while existing automated methods often struggle with imaging artifacts such as condensation, shadows, and contamination.

In this study, we developed the Tuberculosis Microbial Analysis System (TMAS), a machine learning-based tool designed to automate MIC determination from plate images. Training on advanced deep learning models, TMAS can learn to detect bacterial growth with high precision, distinguishing true growth from artefacts such as shadows, bubbles, sediment, condensation and contamination. When tested on a large dataset, TMAS outperformed existing automated methods in accuracy, reliability and efficiency. By reducing the reliance on manual interpretation, TMAS has the potential to streamline TB diagnostics, improve efficiency in laboratories, and improve access to high-quality DST, particularly in resource-limited settings.

## Introduction

Tuberculosis still kills more people each year than any other single pathogen and drug resistance is an increasingly important problem [1]. Whilst the drug development pipeline is better stocked than for decades, the longevity of new drug regimens will depend heavily on our ability to minimise the evolution of resistance by treating patients with drugs to which their tuberculosis is susceptible [2] [3]. Achieving this through the culture-based, phenotypic drug susceptibility testing (pDST) methods in routine use today would be costly and complex [4]. There has consequently been a drive from the WHO and others towards developing and implementing molecular assays, most recently targeted next generation sequencing (tNGS) [5]. However, whilst the molecular determinants of resistance are fairly well understood for many established drugs, there remains much to be learnt for the newly emerging drugs [6].

For the purposes of research and surveillance, an affordable phenotypic assay is therefore required to generate data on a large number of new drugs. 96-well broth microdilution plates are emerging as the most practical approach, and have the added benefit that they generate minimum inhibitory concentrations (MICs) [7]. The experience of the CRyPTIC consortium shows that plate reading can, however, be a challenge. CRyPTIC addressed this by having plates read by laboratory scientists, by a citizen science project, and by a simple image processing tool, and then seeking a consensus from these three methods [8] [9] [10]. Others are now pursuing a similar approach with microtitre plates being used for new drugs in clinical trials, and with a routine plate on its way to commercialisation. Whilst it is likely that experts will continue to be needed for plate reading, a high-quality second reading will be key to ensuring accuracy and efficiency, especially when processing a large number of plates.

Machine learning has demonstrated itself to be well-suited to the interpretation of images, although detecting bacterial growth on microtiter plates poses a challenge. Any method must distinguish between the growth of M. tuberculosis and plate errors such as contamination, condensation, sediment, or air bubbles, whilst not being misled by artefacts such as shadows and water droplets that form on the protective film due to humidity. In this study, the dataset from the CRyPTIC project is utilised to train and test state-of-the-art deep learning models to assess whether this can be achieved. We harness a series of advances in deep learning-based visual object recognition, and object detection specifically [11], to assess its applicability to *M. tuberculosis* pDST.

## Materials and methods

### Dataset

To develop and evaluate the TMAS framework, the publicly available dataset provided by the CRyPTIC Consortium was utilised, which contains extensive phenotypic data of *Mycobacterium tuberculosis* isolates [10]. This dataset comprises:

1. **Image Data:** A total of 15,209 images of 96-well broth microdilution plates captured after 14 (or in a few cases, 21) days of incubation using a *Thermo Fisher Vizion Instrument*. Each image shows an entire plate (referred to as a “plate image”) and includes all 96 wells, each corresponding to different drug concentrations. The plates were set up in two designs:
  - **UKMYC5:** Contains 14 anti-TB drugs and two drug-free positive control wells.
  - **UKMYC6:** An updated design with 13 anti-TB drugs, excluding para-aminosalicylic acid (PAS), and extended concentration ranges for several drugs present in UKMYC5 (Fig 1).
2. **Plate Readings and Ground Truth MICs:** The Minimum Inhibitory Concentration (MIC) values used in this study were derived from the *CRyPTIC* project through a robust consensus process. MIC readings were obtained from three complementary sources: (1) manual readings by expert laboratory technicians using the *Thermo Fisher Vizion* system, which served as a gold-standard reference; (2) automated analyses conducted with *AMyGDA*, a growth detection algorithm designed for reproducible plate image analysis; (3) crowd-sourced evaluations from the *BashTheBug* citizen science project, where non-expert participants analyzed the same images. These readings were integrated by CRyPTIC into a single, standardized consensus MIC value using a purpose-built algorithm [10]. This consensus value served as the ground truth for training and evaluating the performance of TMAS’s growth detection model.

**Fig 1.**
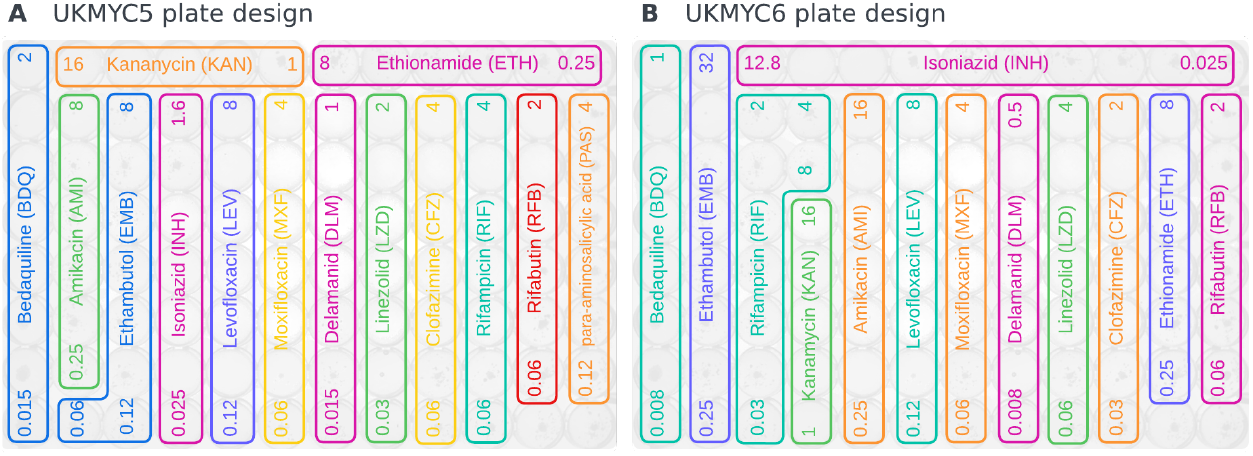
Plate designs for (A) UKMYC5 and (B) UKMYC6. Text labels indicate drug names, while numerical values denote the corresponding drug concentration ranges [7]. Each well has double the drug concentration of the one before. The plates are visualized as background.

**Fig 2.**
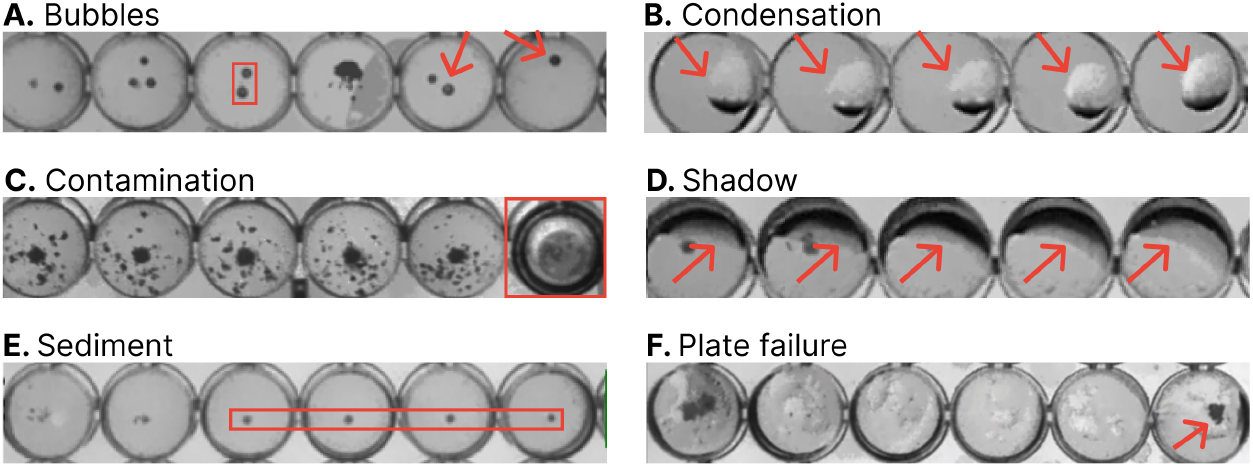
Types of artefacts that can affect MIC determination and model performance, including **(A)** bubbles, **(B)** condensation, **(C)** contamination, **(D)** shadow, **(E)** sediment, and **(F)** plate failure.

### Image Selection and Partitioning

From the complete CRyPTIC dataset of 15,209 plate images, a subset of 4,018 images was randomly selected to train and evaluate TMAS. The selection of a subset of images was to ensure computational efficiency in training and testing the models. The images were chosen randomly to ensure a diverse representation of images, high-quality, low-quality, and challenging edge-case images.

The dataset of 4,018 images for experiments in this work was then partitioned into subsets for training, testing and evaluation purposes, in which the training and validation dataset is 80% and the test set is 20%.

1. **Training and Validation Set:** This set, consisting of 3,215 images (containing 41,850 MIC readings), was used for training and validating the deep learning models for growth detection.
  - **Training Subset:** 75% of the Training and Validation Set (2,411 images or 31,385 MIC readings)
  - **Validation Subset:** 25% of the Training and Validation Set (804 images or 10,465 MIC readings) To mitigate variability introduced by random sampling, the splitting and training process was repeated across five independent runs, with training and validation subsets reshuffled for each run. Model performance for growth detection was evaluated on the validation subsets, and the final results were reported as the mean and standard deviation across the five runs. This approach ensured a more reliable assessment of the growth detection model’s generalizability while reducing potential biases introduced by specific data partitions.
2. **Test Set:** This comprehensive test set, consisting of 803 images (10,454 MIC readings), was used for evaluating TMAS’s performance in predicting MICs.

Additionally, a set of **100 edge-case images**, named the **Edge Case Dataset**, was included to assess TMAS’s robustness under challenging conditions. These images were manually curated from the remaining images in the whole dataset of 15,209 plate images from CryPTIC, and they are characterized by severe artefacts and irregular growth patterns, contributed **1**,**324 MIC readings** to the overall dataset. Together, the Comprehensive Test Set and Edge Case Dataset were used to evaluate TMAS’s MIC prediction performance across diverse conditions and scenarios.

### Source of Errors

In real-life environments, a range of artefacts can significantly impact the quality and clarity of images taken from 96-well plates, potentially affecting model performance. Systematic artefacts that may affect a region or an entire plate include shadows, sediment, and plate failures, which are challenging to detect and avoid. Randomly occurring artefacts, such as bubbles, condensation, or contamination, can lead to nonsensical growth patterns if erroneously detected. These include apparent bacterial proliferation at high antibiotic concentrations—where inhibition was expected—or unexpected absences of growth, causing non-continuous bacterial growth patterns [9]. The *TMAS framework* was specifically designed to mitigate the effects of these artefacts on MIC determination, representing a significant improvement over existing approaches like *AMyGDA*, which are more sensitive to such errors.

### Software

TMAS was built based on advanced deep learning models. It was developed using object-oriented Python3 and implemented with Ultralytics YOLOv8 [12], Keras [13], TensorFlow [14], PyTorch [15] for bacterial growth detection. Image processing techniques were employed using the OpenCV2 Python API [16]. NumPy and Pandas were used to ensure efficient data management. TMAS reports MICs based on the specific plate design and offers the option of saving results in different formats (.csv, .json). TMAS can be installed directly from its PyPi package https://pypi.org/project/tmas using *pip install tmas* or via https://github.com/Oucru-Innovations/tmas.git.

### Solution Design

The TMAS framework includes image preprocessing, selecting the optimal deep learning model for bacterial growth detection, generating and evaluating outputs, and combining growth detection with plate design to report MICs for each drug. The overall architecture is presented in Fig 3.

**Fig 3.**
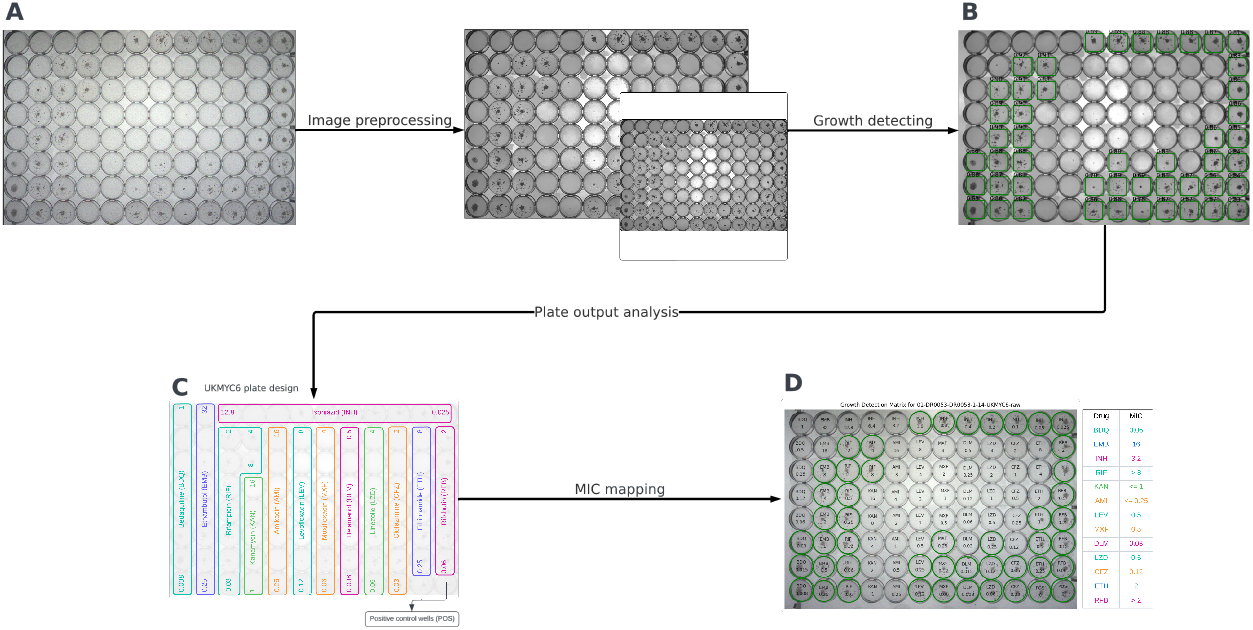
The overall architecture of TMAS. The system consists of four steps to detect growth and determine the MICs of plate images. **(A)** Plate images are preprocessed by applying various image processing techniques. The images are then resized to 512 × 512 pixels and padded to fit the detection model’s input. **(B)** A trained deep learning model is used to detect the bacterial growth in each well of the plates. **(C)** Drug-free, positive control wells are checked for the presence of growth. **(D)** MICs are reported based on growth detection and the corresponding plate design.

Image preprocessing was undertaken to deal with poor image quality, such as mitigating the effects of low contrast and uneven illumination in the acquired images. A mean shift filter [17] was employed to reduce image noise while preserving the integrity of object edges; the Contrast-Limited Adaptive Histogram Equalization (CLAHE) filter [18] was utilised for local contrast enhancement without excessive noise amplification; and pixel histogram normalisation [19] was implemented to optimise overall contrast [9]. We have developed an annotation tool to retrieve consensus MIC values from the CRyPTIC dataset and link plate coordinates to areas of expected microbial growth within the images (see Fig 4). This preparatory step allowed precise labelling of each image and subsequent effective training in later stages of the analysis.

**Fig 4.**
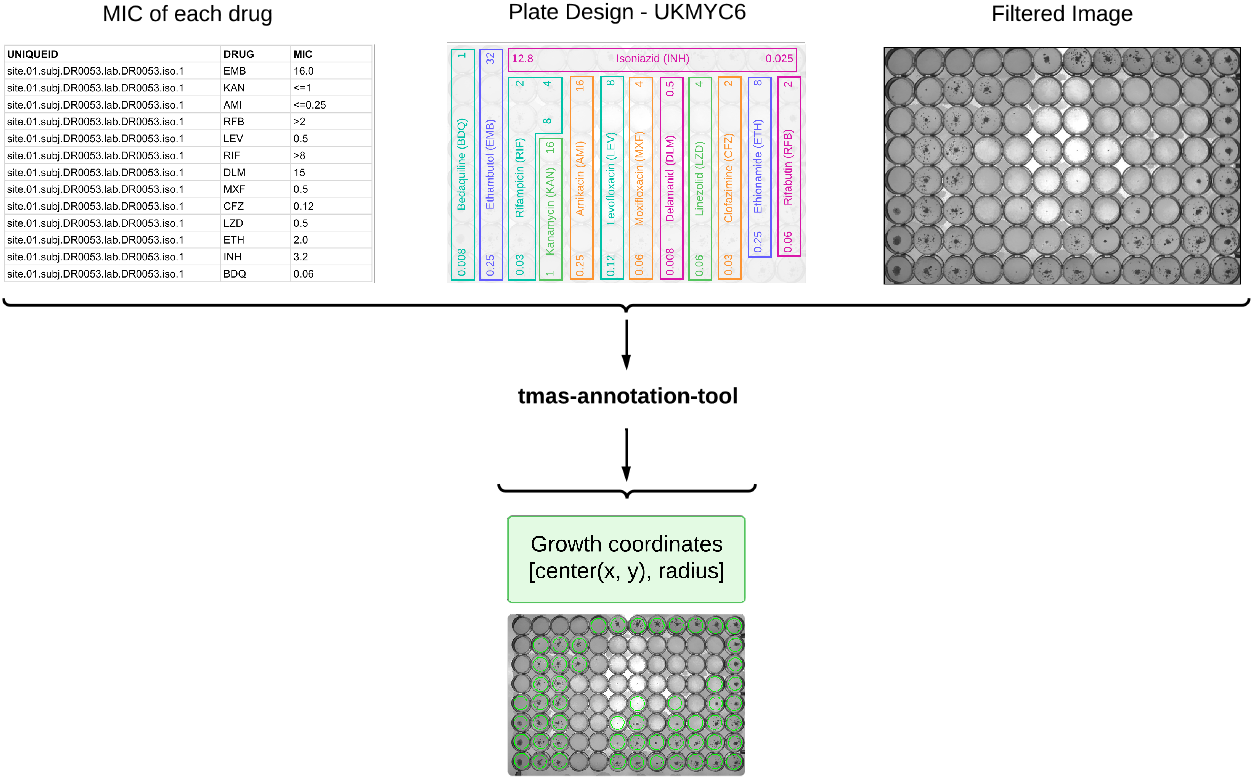
An illustration of annotation tool. The plate images were annotated using the developed tool by retrieving the growth information provided in the plate measurements to determine the growth coordinates in each image, which were then used as bounding boxes for model training.

To standardize the input for the deep learning models, all processed images were resized to 512 × 512 pixels and padded to preserve their unique features while maintaining the original aspect ratio (Fig 3A). The performance of each model was rigorously evaluated using standard metrics — including Precision, Recall, F1 Score, and Mean Average Precision (mAP) for bacterial growth detection — as well as computational efficiency indicators such as throughput.

To ensure the reliability of predictions, a confidence score threshold of 0.5 was applied during the evaluation phase; only detections with confidence scores ≥ 0.5 were considered valid, thereby ensuring a reasonable level of certainty in the results. Following detection, positive control wells outputs were verified (see Fig 3C). Images exhibiting discrepancies or errors were flagged for manual review to further ensure accuracy and robustness.

The Minimum Inhibitory Concentrations (MICs) were defined using a binary growth matrix, where 1 indicates detected bacterial growth and 0 represents no growth. For each drug, if growth was detected across all wells, the MIC was determined to be higher than the highest drug concentration. Conversely, if no growth was observed in any wells, the MIC was recorded as lower or equal to the lowest drug concentration. As part of the output, visualizations of the bacterial growth detection on plate images can be optionally provided alongside the MIC results for all drugs tested (see Fig 3D).

To evaluate TMAS’s performance, we tested it on a comprehensive test set of 803 images, each paired with a corresponding ground truth image. For each drug in a test image, TMAS’s predicted MIC was compared to the ground truth MIC for the same drug. The evaluation focused on the agreement between TMAS predictions and ground truth values based on dilution levels. Essential agreement was defined as the percentage of MIC predictions that were either identical to or within one doubling dilution of the ground truth MIC. Out of 10,454 MIC reading pairs from the 803 plate images, 176 pairs were excluded due to missing ground truth values, resulting in 10,278 MIC reading pairs being analyzed. These comparisons formed the basis for calculating TMAS’s overall essential agreement.

### Experiment Setup

All experiments were conducted on Google Colab using an NVIDIA L4 GPU and local hardware including a B760 Edge Ti WiFi motherboard, a Core i5-12400 CPU, 16 GB DDR5 RAM, an MPG A850GL power supply unit, a MEG S280 cooler, an RTX 4080 SUPRIM graphics card, a M390 500 GB SSD, a GUNGIR 300R case, and peripherals GK30 and GM20. The environment was configured using Python 3.10.12 and PyTorch 2.3.1, utilising CUDA 12.2 (compiler version 12.2.140).

## Results

### Evaluation of TMAS’ Growth Detection Model

Four advanced Deep Learning models (Faster R-CNN, Mask R-CNN, Inception-ResNet, and YOLOv8) were trained and evaluated, focusing on key performance metrics: precision, recall, F1-score, mean average precision (mAP) [20], and throughput (the number of images processed per second, ips). The results, averaged over five runs, are summarized in Table 1.

**Table 1.** Performance comparison of different models, sorted by F1-score in descending order. Ips = images per second.

| Models | Precision | Recall | F1-score | mAP | Throughput |
| --- | --- | --- | --- | --- | --- |
| YOLOv8 | 0.956 ± 0.005 | 0.954 ± 0.004 | 0.955 ± 0.004 | 0.982 ± 0.002 | 69 ips |
| Mask R-CNN | 0.910 ± 0.030 | 0.969 ± 0.019 | 0.938 ± 0.011 | 0.952 ± 0.016 | 19 ips |
| Faster R-CNN | 0.914 ± 0.029 | 0.961 ± 0.017 | 0.937 ± 0.008 | 0.970 ± 0.005 | 25 ips |
| Inception-ResNet | 0.996 ± 0.001 | 0.707 ± 0.083 | 0.826 ± 0.058 | 0.794 ± 0.004 | 15 ips |

Table 1 shows average performance across five runs. YOLOv8 achieved the highest F1-score, mAP, and throughput.

**Table 2.**
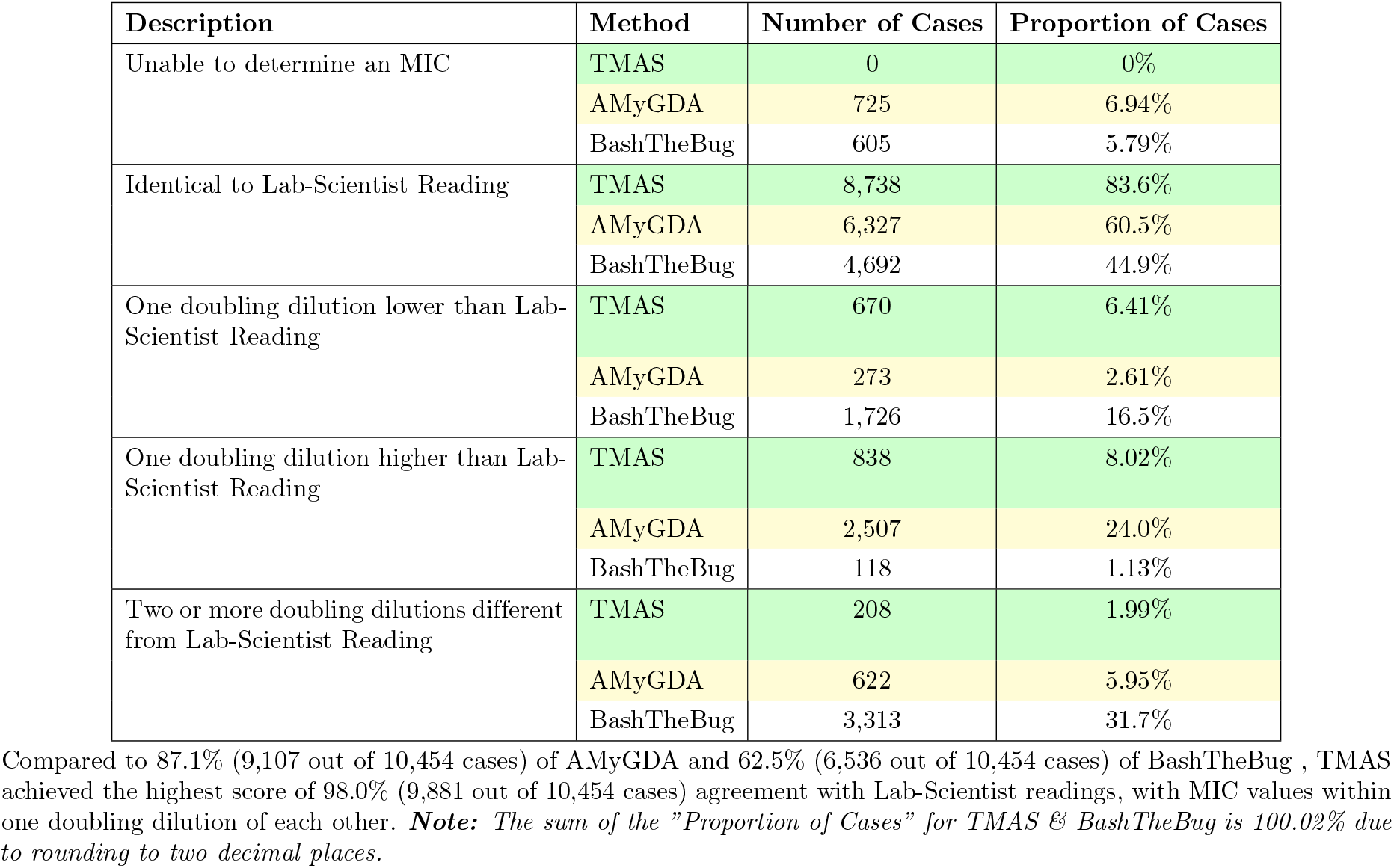
Comparison of MIC values predicted by TMAS, AMyGDA, and BashTheBug against the MIC measured by laboratory scientists on the comprehensive test set.

**Table 3.** Comparison of MIC values measured by TMAS, AMyGDA, and BashTheBug against the MIC measured by laboratory scientists on the edge case dataset.

| Description | Method | Number of Cases | Proportion of Cases |
| --- | --- | --- | --- |
| Unable to determine an MIC | TMAS | 0 | 0% |
|  | AMyGDA | 267 | 20.4% |
|  | BashTheBug | 278 | 21.2% |
| Identical to Lab-Scientist Reading | TMAS | 816 | 62.2% |
|  | AMyGDA | 574 | 43.8% |
|  | BashTheBug | 483 | 36.8% |
| One doubling dilution lower than Lab-Scientist Reading | TMAS | 236 | 18.0% |
|  | AMyGDA | 74 | 5.64% |
|  | BashTheBug | 148 | 11.3% |
| One doubling dilution higher than Lab-Scientist Reading | TMAS | 160 | 12.2% |
|  | AMyGDA | 269 | 20.5% |
|  | BashTheBug | 18 | 1.37% |
| Two or more doubling dilutions different from Lab-Scientist Reading | TMAS | 99 | 7.55% |
|  | AMyGDA | 127 | 9.69% |
|  | BashTheBug | 384 | 29.3% |

TMAS achieved the highest agreement at 92.4% (1,212 out of 1,311 cases), compared to 69.9% (917 out of 1,311 cases) for AMyGDA and 49.5% (649 out of 1,311 cases) for BashTheBug, with agreement defined as MIC values that are identical to or within one doubling dilution of the Lab-Scientist readings. **Note:** Due to rounding to two decimal places, the total proportion of cases sums to 99.95% for TMAS, 99.97% for BashTheBug, and 100.03% for AMyGDA.

Among the evaluated models, YOLOv8 consistently outperformed the others across all metrics, achieving the highest F1-score, mAP, and throughput. Mask R-CNN and Faster R-CNN also demonstrated strong performance, but their F1-scores and throughput were slightly lower than YOLOv8. Inception-ResNet achieved the highest precision but suffered from low recall, which significantly impacted its F1-score and mAP.

Furthermore, an analysis of the learning behaviors of the models, based on loss graphs (provided in the supplementary section), revealed that YOLOv8 exhibited faster convergence and more stable loss reduction during training, indicating superior learning efficiency compared to the other models. This advantage translated into higher detection accuracy and reliability, affirming YOLOv8 as the best-performing model in this study. To further validate YOLOv8, we quantified its inference time on the same testing dataset. With a throughput of 69 images per second, YOLOv8 was the most computationally efficient model, significantly surpassing the throughput of Mask R-CNN (19 ips), Faster R-CNN (25 ips), and Inception-ResNet (15 ips).

Based on these evaluations, YOLOv8 was selected as the optimal model for growth detection and subsequently integrated into TMAS for Minimum Inhibitory Concentration (MIC) extraction. An example of the visualized detection results is shown in Fig 5B, alongside a comparison to the ground truth in Fig 5A.

**Fig 5.**
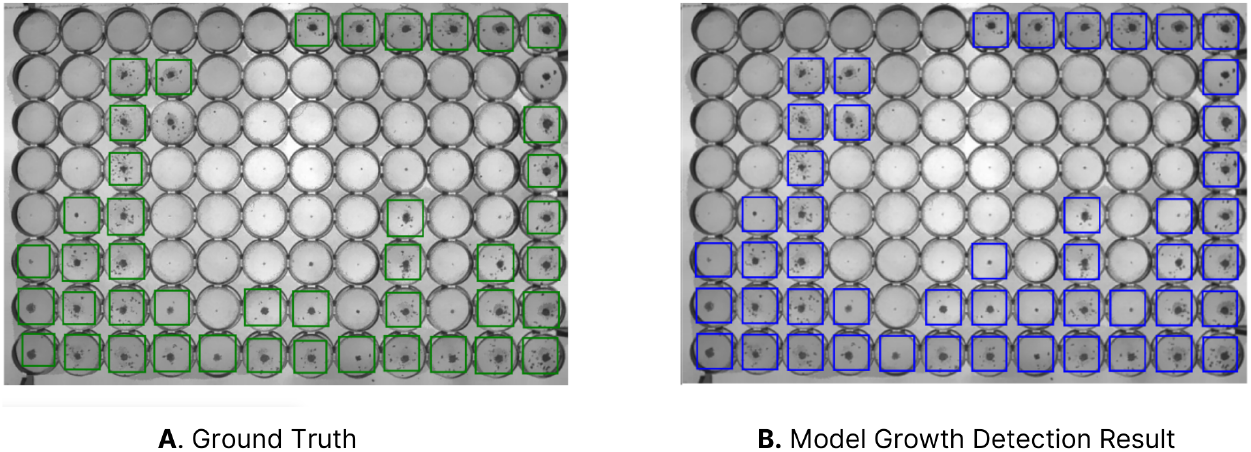
Example of comparison of ground truth and YOLOv8 detection results on a plate image. **(A)** displays ground truth bacterial growth (green bounding boxes), while **(B)** illustrates YOLOv8 predictions (blue bounding boxes).

### TMAS’s MIC Reading Result on Test Set

The TMAS framework was evaluated for its ability to determine Minimum Inhibitory Concentrations (MICs) by integrating output from the bacterial growth detection model applied to 96-well plate images.

The concordance between TMAS’s MIC predictions and the ground truth was high, as demonstrated by a strong linear relationship with an *R*^2^ of 0.94 (Fig 6). To ensure reliability in the essential agreement calculations, 63 additional MIC readings (0.61%) were excluded due to nonsensical growth patterns, such as the detection of bacterial growth in higher concentration wells but not in lower concentration wells. These cases, requiring manual review, were excluded to maintain the validity of the automated evaluation. This resulted in 10,215 MIC readings being used for essential agreement calculations.

**Fig 6.**
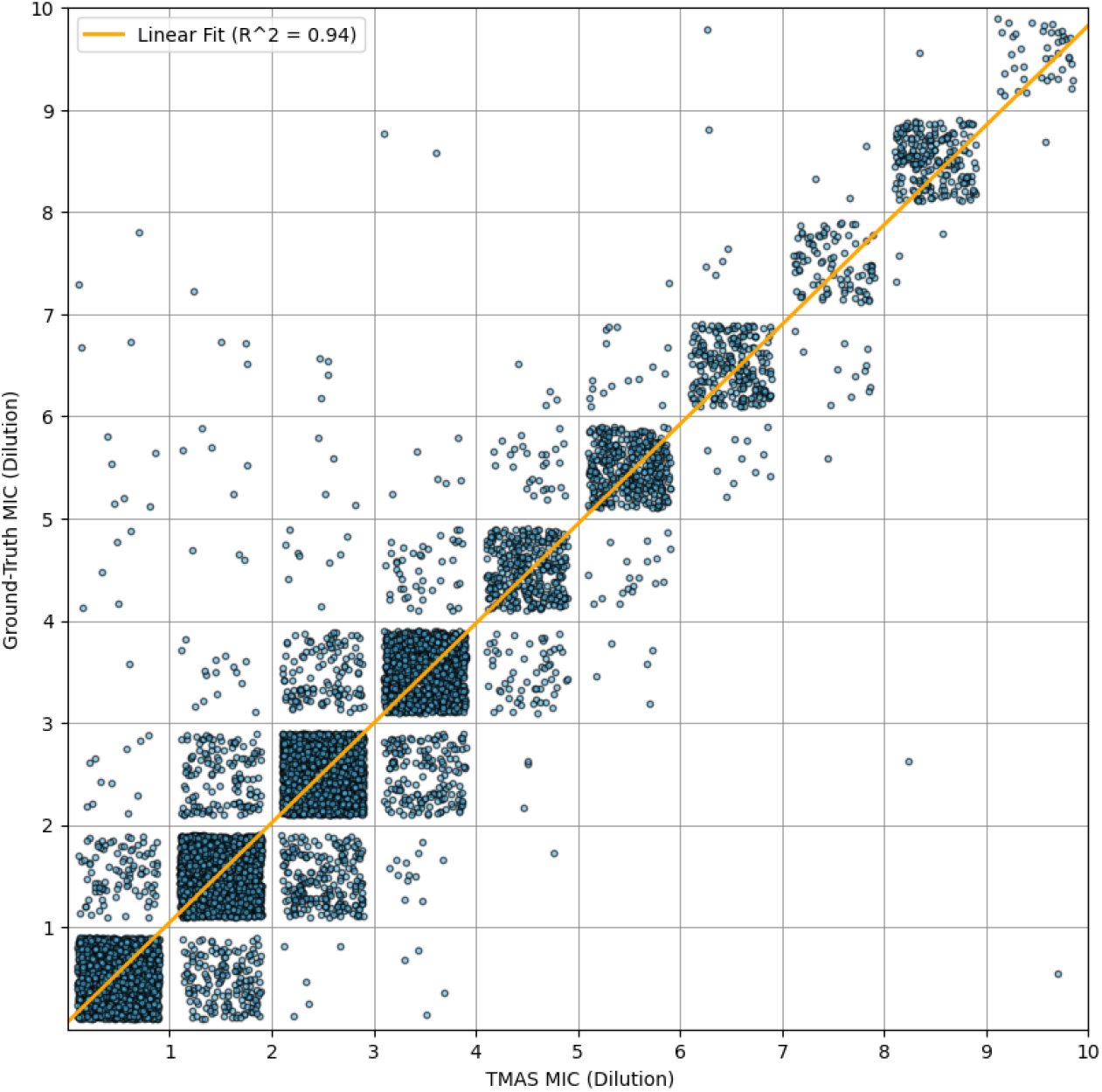
Relationship between TMAS-predicted MIC (Dilution) and ground truth. Each point represents one of the 10,278 pairs of predicted and ground truth MIC dilutions, with a linear regression trend line (*R*^2^ = 0.94) illustrating the strong agreement across all drugs. Variations in the number of wells for each drug may introduce minor biases in the distribution.

TMAS achieved an essential agreement of 98.8% (10,096 out of 10,215 cases) with the ground truth. Of the 10,215 cases, TMAS predictions were identical to the ground truth for 9,009 cases (88.2%), one doubling dilution lower for 645 cases (6.31%), and one doubling dilution higher for 442 cases (4.33%). Predictions that differed by two or more doubling dilutions were classified as errors, accounting for 119 cases (1.16% error rate). This performance significantly exceeds the 90% threshold established by the International Organization for Standardization (ISO) [21], which is commonly used to assess and validate new drug susceptibility testing (DST) methods. These results demonstrate the reliability and accuracy of TMAS in determining MICs under diverse testing conditions.

### TMAS, AMyGDA, and BashTheBug Performance on Test Set compared to Lab-Scientist Readings

To compare TMAS with AMyGDA and BashTheBug, MICs determined by the laboratory-based expert reader were used as the benchmark. This approach provides an independent reference point since our ground truth contains results from AMyGDA and BashTheBug, making it unsuitable for objective evaluation. Each method’s ability to detect bacterial growth was assessed across two test sets: the comprehensive test set and the edge case dataset with plate artefacts.

As observed in Fig 7, the results from TMAS and AMyGDA are relatively similar. Both methods successfully detected limited growth in wells containing EMB, LEV, MXF, LZD, PAS, and KAN. However, the bacterial growth detected by TMAS was consistently reported at least one doubling dilution higher than that observed by laboratory scientists. AMyGDA occasionally failed to detect the same level of growth, resulting in MICs that were one dilution lower than those determined by laboratory scientists. This led to discrepancies in MICs compared to the Lab-Scientist Reading, although results were still within essential agreement. The consensus of the BashTheBug volunteers was less accurate at detecting bacterial growth. In some cases, it failed to identify growth detected by TMAS and AMyGDA, leading to a higher discrepancy in its reported MICs.

**Fig 7.**
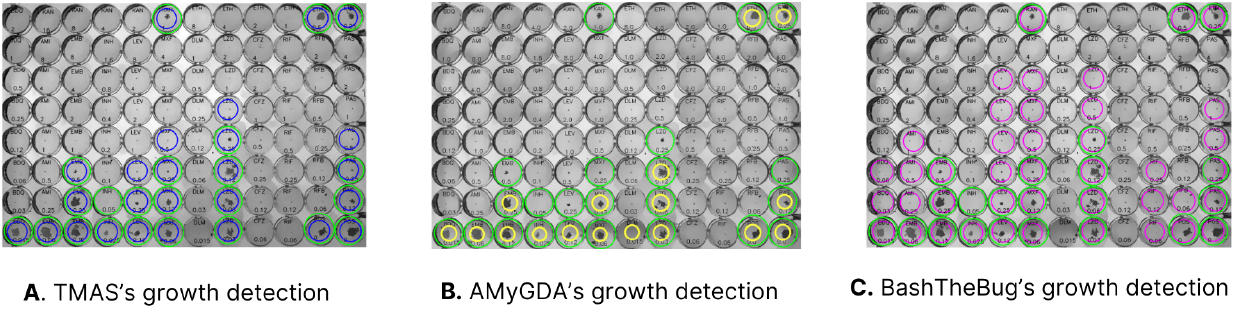
Comparison of TMAS, AMyGDA, and BashTheBug growth detection results. The green circles indicate the Ground Truth data provided by the lab scientists. The left column **(A)** displays the TMAS’s detection results, with bacterial growth marked by blue circles. The middle column **(B)** displays the AMyGDA detection results, with bacterial growth marked by yellow circles. The last column **(C)** represents the BashTheBug detection results, with bacterial growth marked by pink circles.

TMAS achieved the greatest concordance with 83.6% of its readings exactly matching those of lab scientists, compared to 60.5% of AMyGDA’s readings and 44.9% of BashTheBug’s. Moreover, TMAS’s discrepancy rates for readings one doubling dilution off from Lab-Scientist readings stood at 14.43% (6.41% one dilution lower + 8.02% one dilution higher), being lower than AMYGDA’s 26.61% (2.61% + 24.0%) and BashTheBug’s 17.63% (16.5% + 1.13%). AMYGDA’s performance suffers from its reliance on structured scenarios, and it cannot adapt to handle low-quality or distorted images that result from plate artefacts. Only 1.99% of TMAS readings differed by two or more doubling dilutions from Lab-Scientist readings, compared to 5.95% for AMYGDA and 31.7% for BashTheBug.

### TMAS, AMYGDA, BashTheBug Performance on Edge Case Dataset compared to Lab-Scientist Reading

Next we compared the performance of TMAS, AMyGDA, and BashTheBug on hard-to-read plates containing artefacts. Lab-scientist readings again served as the ground truth benchmark for this comparison. 13 out of 1,324 readings from 100 images were excluded due to the lack of ground truth, resulting in 1,311 in total.

As an example in (Fig 8), TMAS (Fig 8A) and BashTheBug (Fig 8C) performed well in detecting bacterial growth among these edge cases. While BashTheBug perfectly aligned with lab-scientist readings in this case, TMAS exhibited one false positive reading for AMI and one false negative reading for CFZ. AMyGDA (Fig 8B) frequently misclassified condensation as bacterial growth, leading to a high rate of incorrect MIC readings. As it relies on crowd-sourced inputs, BashTheBug is particularly susceptible to errors arising from misinterpretations of visual artefacts, such as mistaking sediment for microbial growth or condensation patterns, which increases the risk of inaccuracies in MIC determination.

**Fig 8.**
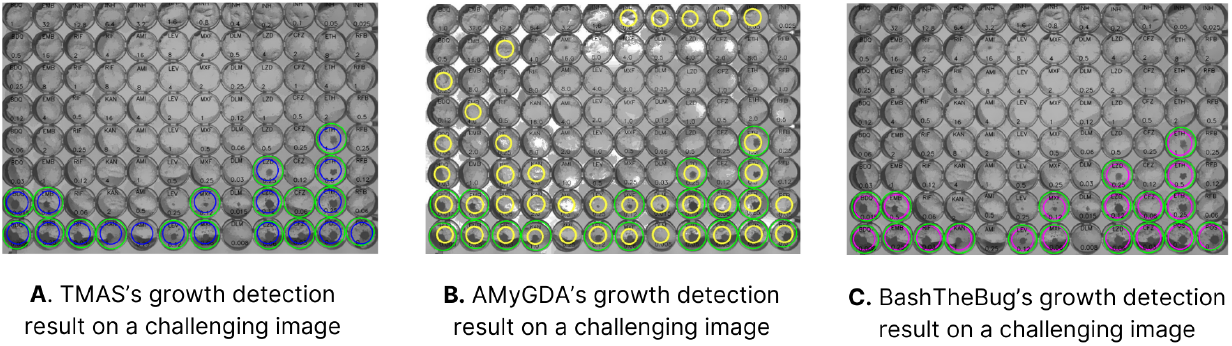
Comparison of TMAS and AMyGDA growth detection results on challenging images. The Ground Truth data provided by the lab scientists is indicated by the green circles. The left-hand plate image **(A)** displays the TMAS’s detection results, with bacterial growth marked by blue circles. The middle plate image displays the AMyGDA detection results, with bacterial growth marked by yellow circles. The right-hand plate image **(C)** represents the BashTheBug detection results, with bacterial growth marked by pink circles.

From the comparison of TMAS, AMyGDA, and BashTheBug on the edge case test set, we can see that TMAS consistently demonstrated the closest correlation with the Lab-Scientist Readings, achieving a match in 62.2% of cases, compared to 43.8% of AMyGDA’s readings and 36.8% of BashTheBug’s. It also demonstrated the least discrepancies, with only 7.55% of cases differing by two or more dilutions, compared to 9.69% for AMyGDA and 29.3% for BashTheBug. Additionally, TMAS achieved the lowest error rate, with no MICs undetermined, compared to error rates of 20.4% for AMyGDA and 21.2% for BashTheBug. TMAS is therefore more accurate, reliable, and robust than existing methods, even when presented with challenging edge cases.

## Discussion

We have developed and evaluated the Tuberculosis Microbial Analysis System (TMAS), an automated framework for detecting bacterial growth from images of 96-well microtiter plates and determining Minimum Inhibitory Concentrations (MICs). Microtiter plates are an efficient means of simultaneously measuring MICs for multiple drugs, and cost less than traditional DST methods, such as the Mycobacterial Growth Indicator Tube (MGIT), would if used to assay MICs for this many drugs. TMAS leverages advanced deep learning algorithms to accurately read 96-well broth microdilution plates. On the comprehensive test set, TMAS achieved an essential agreement with the ground truth of 98.8%, significantly exceeding the 90% minimum required for drug susceptibility testing methods by ISO 20776-2:2021 [21]. Furthermore, TMAS was able to interpret challenging images, maintaining high agreement rates despite the presence of artefacts and low-quality images. It also achieved a trade-off between computational efficiency and accuracy, with an average inference time of 14.3 milliseconds per image and a throughput of 69 images per second, ensuring its practical applicability.

TMAS addresses several critical limitations of earlier systems such as AMyGDA and BashTheBug, offering notable improvements in accuracy, robustness, and reliability. Our evaluations demonstrated that TMAS consistently outperformed AMyGDA and BashTheBug across multiple metrics. On the comprehensive test set, TMAS achieved 83.6% agreement with lab-scientist readings, compared to 60.5% for AMyGDA and 44.9% for BashTheBug. Additionally, TMAS reported discrepancies (one doubling dilution off) in only 14.4% of cases, lower than AMyGDA’s 26.6% and BashTheBug’s 17.6%. TMAS also exhibited the lowest error rate (1.99%) for readings that were two or more dilutions different from the ground truth, compared to 5.95% for AMyGDA and 31.7% for BashTheBug.

BashTheBug, a crowd-sourced citizen science platform, exhibited variability in performance due to its reliance on non-expert contributors. While the “wisdom of the crowd” approach helps average out individual errors, the project’s high Gini coefficient indicates an uneven distribution of classifications, with a small subset of volunteers contributing disproportionately more than others [8]. However, as each volunteer could only contribute a single measurement per MIC, the overall impact of this imbalance was limited. A more critical limitation of BashTheBug was that volunteers tended to be conservative in their classifications, often overcalling bacterial growth as a precautionary measure. This tendency also increased the likelihood of misclassifying contamination or other artefacts as bacterial growth, leading to inflated MIC values.

Similarly, AMyGDA, an automated image-processing-based tool, exhibited some limitations due to its reliance on simple techniques [9]. AMyGDA primarily detects bacterial growth by counting dark pixels in the center of wells, rendering it highly susceptible to artefacts. Unlike deep learning-based approaches, which can learn image patterns to distinguish between true bacterial growth and artefacts, AMyGDA requires manual parameter tuning to mitigate false positives. This often results in undercalling MICs compared to laboratory scientists’ readings, as a conservative threshold is applied to minimize erroneous classifications. Although AMyGDA employs mean shift filtering (MSF) and contrast-limited adaptive histogram equalization (CLAHE)—techniques also utilized in TMAS for preprocessing—these methods alone are insufficient to accurately detect bacterial growth in low-quality images. By contrast, TMAS’s deep learning-based approach leverages learned feature representations to robustly differentiate growth from artefacts across diverse imaging conditions, enhancing both accuracy and reliability.

TMAS demonstrated its robustness when evaluated on an edge case dataset characterized by severe artefacts and irregular growth patterns. TMAS achieved 62.2% agreement with lab-scientist readings, outperforming AMyGDA (43.8%) and BashTheBug (36.8%). Additionally, TMAS reported discrepancies (one doubling dilution off) in 31.6% of cases, compared to AMyGDA’s 28.6% and BashTheBug’s 22.7%. TMAS also exhibited the lowest rate of major discrepancies, with only 7.55% of cases differing by two or more dilutions, compared to 9.69% for AMyGDA and 29.3% for BashTheBug. These results emphasize TMAS’s ability to minimize prediction errors and maintain high performance under various conditions, making it a reliable tool for applications in resource-limited settings.

In principle, TMAS could be adapted for real-time monitoring of bacterial growth. For example, laboratories could capture images at regular intervals (e.g., every six hours) following seven days of incubation, enabling TMAS to detect changes in bacterial growth and report MICs “when ready.” This approach may yield valuable insights into growth kinetics and accelerate clinical decision-making. However, such a protocol would require repeatedly removing plates from the incubator—often relying on a single, expensive Thermo Fisher Vizion instrument—thereby increasing labor demands, contamination risk, and overall complexity. Further, the Vizion hardware itself may be inaccessible to laboratories with limited resources. Although a mobile solution might circumvent the need for specialized equipment, it would demand extensive model optimization and a mobile app interface, along with potential costs for cloud-based inference if on-device processing proves infeasible.

Several aspects to consider around developing and using TMAS. First, while TMAS was developed for UKMYC5 and UKMYC6 plate layouts, it could potentially function on modified plates with additional wells or altered drug ranges because the main task remains to detect growth. However, achieving optimal accuracy on substantially altered designs may require adjusting the annotation process, updating the plate design map within TMAS, and potentially retraining certain aspects of the model. Second, TMAS currently produces a binary (growth vs. no growth) classification within each well based on bounding boxes for the growth detection phase. Although this approach effectively localizes bacterial growth, implementing additional post-processing to quantify partial or percentage growth could offer further granularity for research or clinical purposes. Finally, adapting TMAS to novel platforms—such as mobile devices or alternative imaging setups—would require further development to ensure consistent model performance, sufficient computational capacity, and stable lighting conditions.

A notable strength of TMAS is its demonstrated ability to generalize across diverse conditions, as evidenced by its performance on both a broad comprehensive test set and a curated “edge-case” dataset. The edge-case dataset, characterized by severe artefacts and irregular growth patterns, further challenged the model beyond routine scenarios. Future developments could further enhance TMAS’s capabilities. These include gathering images from multiple imaging systems aside from the Thermo Fisher Vizion instrument, refining the detection pipeline to account for partial or percentage growth, and exploring integrated incubator–camera workflows that automate imaging. Additionally, customizing the model’s loss function to accommodate varying phenotype quality (e.g., high, medium, or low) could improve its predictive accuracy in heterogeneous datasets.

TMAS is a significant advancement in the automated, high-throughput reading of broth microdilution plates. It outperformed existing models in terms of accuracy and reliability, improving on the limitations of earlier approaches like AMyGDA and BashTheBug. Future enhancements, such as real-time monitoring and hybrid approaches, could extend its utility, making it a valuable tool for clinical microbiology laboratories and research institutions tackling multidrug-resistant tuberculosis.

## Supporting information

**S1 Fig. Loss graph for different proposed models**. Analysis of the training and validation loss graphs for each state-of-the-art model, including YOLOv8, FasterRCNN, MaskRCNN, Inception-ResNet, represented by blue and orange lines, respectively.

## Acknowledgments

We would like to express our gratitude to NVIDIA and MSI for providing powerful components that greatly assisted our experiments with various state-of-the-art deep learning models. Additionally, we extend our sincere thanks to RMIT University for providing the budget that enabled us to utilize cloud services for experimenting with our framework, which was essential for our research. TMW is a Wellcome Trust Clinical Career Development Fellow (214560/Z/18/Z). Funded by Wellcome (106680/B/14/Z). PWF would like to acknowledge funding from the National Institute for Health Research (NIHR) Health Protection Research Unit (HPRU) in Healthcare Associated Infections and Antimicrobial Resistance at Oxford University in partnership with UK Health Security Agency [HPRU-2012-10041], the National Institute for Health Research (NIHR) and Oxford Biomedical Research Centre (BRC). For the purpose of open access, the author has applied a CC BY public copyright licence to any Author Accepted Manuscript version arising from this submission. The findings and conclusions in this report are solely the responsibility of the authors and do not necessarily represent the official views of the NHS, the NIHR or the Department of Health and Social Care.

## Author Contributions

**Conceptualization:** Timothy M Walker, Philip W Fowler, Hai Ho Bich.

**Formal Analysis:** Hoang-Anh T Vo, Sang Nguyen, Han Nguyen.

**Funding Acquisition:** Thuy Thi Nguyen, Timothy M Walker, Philip W Fowler, Ai- Quynh T Tran.

**Methodology:** Hoang-Anh T Vo.

**Project Administration:** Thuy Thi Nguyen, Hoang-Anh T Vo.

**Resources:** Philip W Fowler, Timothy M Walker.

**Software:** Hoang-Anh T Vo, Sang Nguyen, Han Nguyen, Ai-Quynh T Tran.

**Supervision:** Timothy M Walker, Thuy Thi Nguyen, Hai Ho Bich.

**Validation:** Thuy Thi Nguyen, Timothy M Walker, Hoang-Anh T Vo.

**Visualization:** Hoang-Anh T Vo, Sang Nguyen, Ai-Quynh T Tran, Han Nguyen.

**Writing – Original Draft:** Hoang-Anh T Vo, Sang Nguyen, Ai-Quynh T Tran, Han Nguyen, Timothy M Walker.

**Writing – Review & Editing:** Hoang-Anh T Vo, Sang Nguyen, Ai-Quynh T Tran, Han Nguyen, Philip W Fowler, Timothy M Walker, Thuy Thi Nguyen.

## References

[1] World Health Organization. Global Tuberculosis Report 2023. Geneva: World Health Organization, 2023. isbn: 9789240083851. url: https://www.who.int/publications/i/item/9789240083851.

[2] Véronique A. Dartois and Eric J. Rubin. “Anti-tuberculosis treatment strategies and drug development: challenges and priorities”. en. In: Nature Reviews Microbiology 20.11 (Nov. 2022), pp. 685–701. issn: 1740-1526, 1740-1534. doi: 10.1038/s41579-022-00731-y.

[3] Annelies Van Rie et al. “Balancing access to BPaLM regimens and risk of resistance”. en. In: The Lancet Infectious Diseases 22.10 (Oct. 2022), pp. 1411–1412. issn: 14733099. doi: 10.1016/S1473-3099(22)00543-6.

[4] Paola M. V. Rancoita et al. “Validating a 14-Drug Microtiter Plate Containing Bedaquiline and Delamanid for Large-Scale Research Susceptibility Testing of Mycobacterium tuberculosis”. en. In: Antimicrobial Agents and Chemotherapy 62.9 (Sept. 2018), e00344–18. issn: 0066-4804, 1098-6596. doi: 10.1128/AAC.00344-18.

[5] WHO Global Tuberculosis Programme (GTB). Use of targeted next-generation sequencing to detect drug-resistant tuberculosis: rapid communication, July 2023. url: https://www.who.int/publications/i/item/9789240076372.

[6] Timothy M Walker et al. “The 2021 WHO catalogue of Mycobacterium tuberculosis complex mutations associated with drug resistance: a genotypic analysis”. en. In: The Lancet Microbe 3.4 (Apr. 2022), e265–e273. issn: 26665247. doi: 10.1016/S2666-5247(21)00301-3.

[7] The CRyPTIC Consortium et al. Epidemiological cutoff values for a 96-well broth microdilution plate for high-throughput research antibiotic susceptibility testing of M. tuberculosis. en. Mar. 2021. doi: 10.1101/2021.02.24.21252386.

[8] Philip W Fowler et al. “A crowd of BashTheBug volunteers reproducibly and accurately measure the minimum inhibitory concentrations of 13 antitubercular drugs from photographs of 96-well broth microdilution plates”. en. In: eLife 11 (May 2022), e75046. issn: 2050-084X. doi: 10.7554/eLife.75046.

[9] Philip W. Fowler et al. “Automated detection of bacterial growth on 96-well plates for high-throughput drug susceptibility testing of Mycobacterium tuberculosis”. en. In: Microbiology 164.12 (Dec. 2018), pp. 1522–1530. issn: 1350-0872, 1465-2080. doi: 10.1099/mic.0.000733.

[10] The CRyPTIC Consortium. “A data compendium associating the genomes of 12,289 Mycobacterium tuberculosis isolates with quantitative resistance phenotypes to 13 antibiotics”. en. In: PLOS Biology 20.8 (Aug. 2022). Ed. by Jason Ladner, e3001721. issn: 1545-7885. doi: 10.1371/journal.pbio.3001721.

[11] Iqbal H. Sarker. “Deep Learning: A Comprehensive Overview on Techniques, Taxonomy, Applications and Research Directions”. en. In: SN Computer Science 2.6 (Nov. 2021), p. 420. issn: 2662-995X, 2661-8907. doi: 10.1007/s42979-021-00815-1.

[12] Ultralytics: YOLO Vision. en. 2024. url: https://github.com/ultralytics/ultralytics (visited on 06/01/2024).

[13] François Chollet. Deep Learning with Python, Second Edition. eng. New York: Manning Publications Co. LLC, 2021. isbn: 978-1-61729-686-4 978-1-63835-009-5.

[14] Martín Abadi et al. TensorFlow: Large-Scale Machine Learning on Heterogeneous Distributed Systems. 1603.04467 [cs]. Mar. 2016. doi: 10.48550/arXiv.1603.04467.

[15] Adam Paszke et al. PyTorch: An Imperative Style, High-Performance Deep Learning Library. 1912.01703 [cs]. Dec. 2019. doi: 10.48550/arXiv.1912.01703.

[16] Alex Zelinsky. “Learning OpenCV—Computer Vision with the OpenCV Library (Bradski, G.R. et al.; 2008)[On the Shelf]”. In: IEEE Robotics & Automation Magazine 16.3 (Sept. 2009), pp. 100–100. issn: 1070-9932. doi: 10.1109/MRA.2009.933612.

[17] D. Comaniciu and P. Meer. “Mean shift: a robust approach toward feature space analysis”. In: IEEE Transactions on Pattern Analysis and Machine Intelligence 24.5 (May 2002), pp. 603–619. issn: 01628828. doi: 10.1109/34.1000236.

[18] Karel J Zuiderveld and others. “Contrast limited adaptive histogram equalization.” In: Graphics gems 4.1 (1994). Publisher: Academic Press, Boston, MA, USA, pp. 474–485.

[19] Rafael Gonzalez and Zahraa Faisal. Digital Image Processing Second Edition. Prentice Hall, June 2019.

[20] Richard Szeliski. Computer vision: algorithms and applications. Springer Nature, 2022.

[21] International Organization for Standardization. ISO 20776-2:2021 Clinical laboratory testing and in vitro diagnostic test systems — Susceptibility testing of infectious agents and evaluation of performance of antimicrobial susceptibility test devices Part 2: Evaluation of performance of antimicrobial susceptibility test devices against reference broth micro-dilution. Tech. rep. ISO 20776-2:2021. Geneva, Switzerland.: ISO, 2021.

